# HIV-1 Nefs are cargo-sensitive AP-1 trimerization switches in tetherin and MHC-I downregulation

**DOI:** 10.1101/276733

**Authors:** Kyle L. Morris, Cosmo Z. Buffalo, Christina M. Stürzel, Elena Heusinger, Frank Kirchhoff, Xuefeng Ren, James H. Hurley

## Abstract

The HIV accessory protein Nef counteracts immune defenses by subverting coated vesicle pathways. The 3.7 Å cryo-EM structure of a closed trimer of the clathrin adaptor AP-1, the small GTPase Arf1, HIV-1 Nef, and the cytosolic tail of the restriction factor tetherin suggested a mechanism for inactivating tetherin by Golgi retention. The 4.3 Å structure of a mutant Nef-induced dimer of AP-1 showed how the closed trimer is regulated by the dileucine loop of Nef. HDX-MS and mutational analysis were used to show how cargo dynamics leads to alternative Arf1 trimerization, directing Nef targets to be either retained at the trans-Golgi or sorted to lysosomes. Phosphorylation of the NL4-3 M-Nef was shown to regulate AP-1 trimerization, explaining how O-Nefs lacking this phosphosite counteract tetherin but most M-Nefs do not. These observations show how the higher-order organization of a vesicular coat can be allosterically modulated to direct cargoes to distinct fates.

---

The primate lentiviruses HIV and SIV use their accessory proteins Nef and Vpu to remodel the protein repertoire at the surface of infected cells (Kirchhoff, 2010; Tokarev and Guatelli, 2011). This is the primary means whereby these viruses circumvent host defenses to evade immune destruction of infected cells and promote the release of fully infectious particles. MHC-I (Schwartz et al., 1996), CD4 (Garcia and Miller, 1991), SERINC3/5 (Rosa et al., 2015; Usami et al., 2015), and tetherin (Neil et al., 2008; Waheed et al., 2008) are among the major proteins removed from the cell surface in primate lentiviral infection. These proteins are kept away from the surface by various mechanisms, some conserved, and others variable due to adaptations to different primate hosts.

For all primate lentiviruses, MHC-I is targeted at the *trans*-Golgi network (TGN) via the hijacking of clathrin, the clathrin adaptor complex AP-1, and the Golgi-specific small G-protein Arf1 (Coleman et al., 2006; Roeth et al., 2004). In T-cells, Nef typically directs MHC-I into post-Golgi clathrin-coated vesicles (CCVs) that ultimately sort MHC-I to the lysosome, where it is degraded (Roeth et al., 2004; Schwartz et al., 1996). In other cell types, HIV-1 Nef accumulates with MHC-I at the TGN (Greenberg et al., 1998; Le Gall et al., 1998) or endosomes (Lubben et al., 2007). The different fates of MHC-I appear to be a function of phosphoregulation (Kasper et al., 2005). Despite the different outcomes, AP-1 is required for both pathways. Thus, Nef and AP-1 engagement can have different consequences when presented with different cargo signals or when some part of the system is differentially phosphorylated.

Tetherin, which prevents the release of progeny virions from the cell surface, experiences an even wider variety of fates in various lentiviral infections. Pandemic group M HIV-1 strains target tetherin for downregulation by Vpu (Neil et al., 2008; Van Damme et al., 2008), which can either drive TGN accumulation (Schmidt et al., 2011) or lysosomal degradation of tetherin (Kueck and Neil, 2012). AP-1 hijacking by Vpu at the TGN appears to be involved in this process (Jia et al., 2014; Kueck et al., 2015). Most SIVs, however, use Nef for tetherin downregulation (Sauter et al., 2009; Zhang et al., 2009), usually by hijacking the plasma membrane clathrin adaptor complex AP-2 and promoting internalization of tetherin from the plasma membrane (Serra-Moreno et al., 2013; Zhang et al., 2011) and its subsequent lysosomal degradation. HIV-1 O group Nefs (Kluge et al., 2014), and a few M-Nefs (Arias et al., 2016), inactivate tetherin by driving its accumulation at the TGN, presumably by the TGN-specific clathrin adaptor AP-1. The acquisition of distinct tetherin inactivation mechanisms is considered a major aspect of HIV-1 evolution and the pandemic spread of HIV/AIDS (Sauter et al., 2010). Yet the structural and biochemical mechanisms underlying differential inactivation by TGN accumulation or lysosomal degradation are currently unknown.

AP-1 is the heterotetrameric clathrin adaptor complex devoted to the biogenesis of CCVs at the TGN. AP-1 binds to membranes and transmembrane protein cargo on one face, and to clathrin on the other, thereby generating cargo-loaded CCVs (Traub and Bonifacino, 2013). AP-1 consists of two large subunits, β1 and γ, a medium subunit, µ1, and a small subunit σ1 (Traub and Bonifacino, 2013). The large subunits consist of ∼600-residue helical solenoids followed by a linker and an appendage domain. β1 contains the clathrin-binding domain, and both β1 and γ bind to the small G-protein Arf1 (Ren et al., 2013). AP-1 binds to cargo bearing either YxxΦ (where Φ = hydrophobic) or (D/E)XXXLL (LL for short) motifs. YxxΦ motifs bind to the C-terminal domain (CTD) of µ1 (Owen and Evans, 1998), while LL motifs bind to a hemicomplex formed by σ1 and γ (Doray et al., 2007; Jackson et al., 2010; Traub and Bonifacino, 2013). In its inactive cytosolic state, the helical solenoids block the YxxΦ and LL cargo sites on µ1 and σ1-γ, a conformation referred to as the locked state (Collins et al., 2002; Heldwein et al., 2004; Jackson et al., 2010). Arf1 triggers the recruitment of AP-1 to the TGN membrane, where it has further profound effects on AP-1 structure, driving both its allosteric unlocking and its dimerization (Ren et al., 2013).

HIV-1 Nef targets Arf1-unlocked AP-1 via both of its two cargo sites. The cytosolic tail of MHC-I contains an incomplete YxxΦ signal, which is not competent to bind AP-1 on its own. Nef forms a ternary complex that stabilizes the binding of the MHC-I tail to the YxxΦ binding site of µ1 (Jia et al., 2012). Multiple regions of Nef contribute the binding site, with the Pro-rich region 72-78 most prominent among them. Nef can also hijack the LL binding site, as illustrated by the structure of Nef bound to the α-σ2 hemicomplex of AP-2 (Ren et al., 2014), which is well conserved with the γ-σ1 complex of AP-1. Vpu targets the LL site on AP-1 in a similar manner, promoting AP-1 unlocking and making the µ1 CTD available to bind a variant (Shen et al., 2015) motif in the tetherin tail (Jia et al., 2014). In the presence of Arf1 and cargo, Nef induces the oligomerization of unlocked AP-1 into two types of trimers (Shen et al., 2015). One type of trimer ("open") is capable of oligomerizing into an extended hexagonal lattice whose symmetry and dimensions match those of clathrin, and this appears to serve as a mechanism to accelerate CCVs formation (Shen et al., 2015). The other type of trimer ("closed") does not have the same lattice forming property. Its function has been unclear. Yet this state is observed reproducibly in the presence of the tetherin cargo signal.

In this study, we sought to determine whether differential trimer formation could account for differential cargo fate. We determined the structure of the tetherin-bound closed trimer at 3.7 Å resolution by cryo-electron microscopy. The structure showed that Nef uniquely stabilizes the closed trimer by bridging the YxxΦ of one AP-1 to the LL of another. We found that the Nef LL motif is critical to its ability to induce closed trimer formation. Using HDX-MS to probe protein dynamics, we found that the competence of the Nef LL to bridge between AP-1 copies depends on the dynamics of the YxxΦ-µ1 interaction, explaining why tetherin promotes closed trimer formation and MHC-I does not. Analysis of these structures and a related structure of an Arf1-mediated trimer of COPI (Dodonova et al., 2017; Dodonova et al., 2015) led to a model for cargo-regulated alternative trimer formation via the differential bridging and differential use of the Arf1 binding sites of β1 and γ. We also showed that the M-Nef of HIV-1 NL4-3, but not a paradigmatic O-Nef, is subject to phosphoinactivation of dileucine loop bridging and closed trimer formation. This model explains why most M-Nefs are not capable of effectively counteracting human tetherin. We went onto verify this model functionally, providing a structural mechanism to explain how Nef can target the same AP-1:Arf1 system to direct cargoes to different fates.

## Results

### Atomistic structure of the closed AP-1 trimer

The complex of NL4-3 Nef fused to the cytosolic tail of human tetherin was reconstituted with Arf1-GTP and the heterotetrameric core of AP-1. The complex was purified by size exclusion chromatography (SEC), the peak corresponding to a trimer of AP-1 complexes was isolated, and the sample subjected to cryo-EM (Figure S1-3). Both AP-1 trimers and monomers were identified, and their structures reconstructed (Figure 1A-B and Figure S1, 2D, 3). The AP-1 monomer resembles the previously determined hyper-unlocked conformation determined in the presence of a tetherin-Vpu fusion (Jia et al., 2014), with the addition of Arf1 molecules bound to both the β1 and γ sites (Figure S2D). The AP-1 trimer consists of hyper-unlocked monomers mutually arranged in the "closed" conformation previously described at 7 Å resolution (Shen et al., 2015). The structure is non-C3 symmetric (Figure 1C) and was resolved to 3.73 Å (Figure S1A, 2A-B, 3A-F), revealing many new aspects of its organization and Nef, Arf1, and cargo interactions.

**Figure 1.**
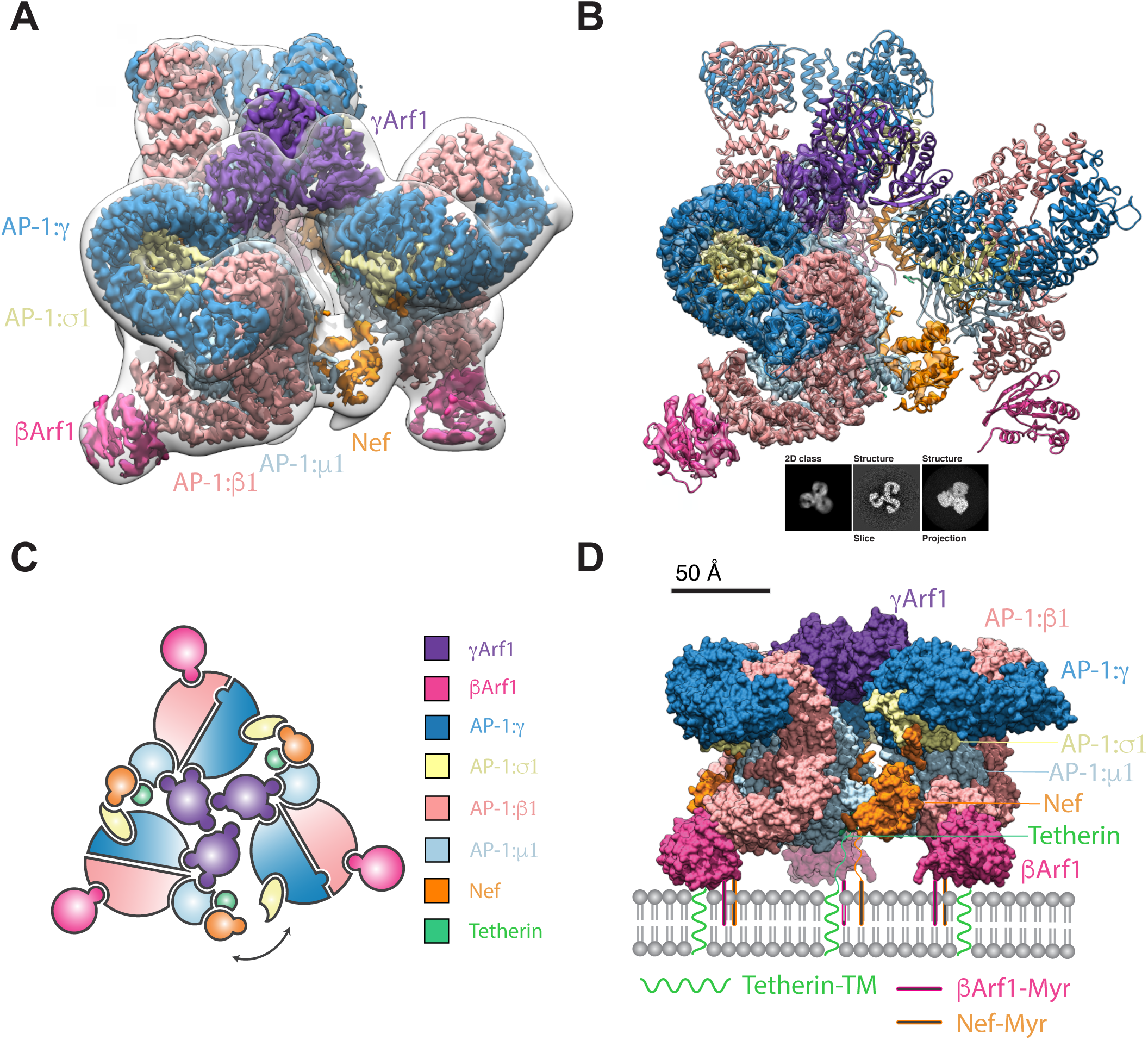
Atomistic structure of the AP1:Arf1:tetherin-Nef trimer. (A) The closed trimeric state consists of three copies of γArf1 assembling three AP-1 cores with one copy of HIV-Nef at the AP1 dimer interface. The subunit reconstruction is shown in the context of the transparent whole trimer reconstruction. The AP-1 core and γArf1 subunits are sharpened at -50 Å^2^, Nef and βArf1 subunits are unsharpened and the whole closed trimer low pass filtered to 18 Å for clarity. (B) The trimer subunit contains the AP-1 core, two copies of Arf1 and one copy of Tetherin-HIV-Nef. A 2D class and corresponding volume slice and projection are shown for reference. (C) A subunit schematic for the closed trimer showing interaction interfaces and the break from perfect three-fold symmetry of the complex allowing movement of the cargo Nef bridge. (D) A model for the closed trimer docked at the trans golgi network. See also Figure S1-3.

The overall trimer is approximately a prism with triangular edges of ∼190 Å and a height of ∼110 Å. AP-1 molecules do not make direct contact with one another. Instead, the trimer is bridged at the top (in the view shown in Figure 1A) by a trimer of Arf1 molecules bound to γ1 (hereafter, "γArf1"), and at the bottom, by Nef. Another three Arf1 molecules are bound to β (hereafter, “βArf1”). These molecules are at the periphery and are not involved in inter-AP-1 bridges.

Two of the three inter-AP-1 gaps at the bottom of the assembly are ∼30 Å across and spanned by ordered Nef molecules (Figure 2A-E), while the third opening, at ∼40 Å, is too wide to be spanned. The inability of Nef to simultaneously span all three gaps accounts for the non-C3 symmetric nature of the assembly. Nef makes extensive contacts with the µ1-CTD and the tetherin tail, which resemble contacts (Figure 2A-B) described for the µ1-CTD:Nef:MHC-I tail complex (Jia et al., 2012). The Nef LL motif is well-defined in density and engages the γ-σ1 (Figure 2C) in trans with respect to the µ1 interaction. The Nef sequences 149-156 and 167-179 are disordered over gaps of 14 and 21 Å, respectively between the LL motif and the Nef core. The Nef core loop consisting of residues 92-94 could make electrostatic contact with the N-terminus of β1, in cis with respect to the LL binding site. Collectively, these observations show how Nef promotes the closed trimer assembly of AP-1 by bridging *in trans* between the µ1-CTD and LL sites of two pairs of AP-1 monomers.

**Figure 2.**
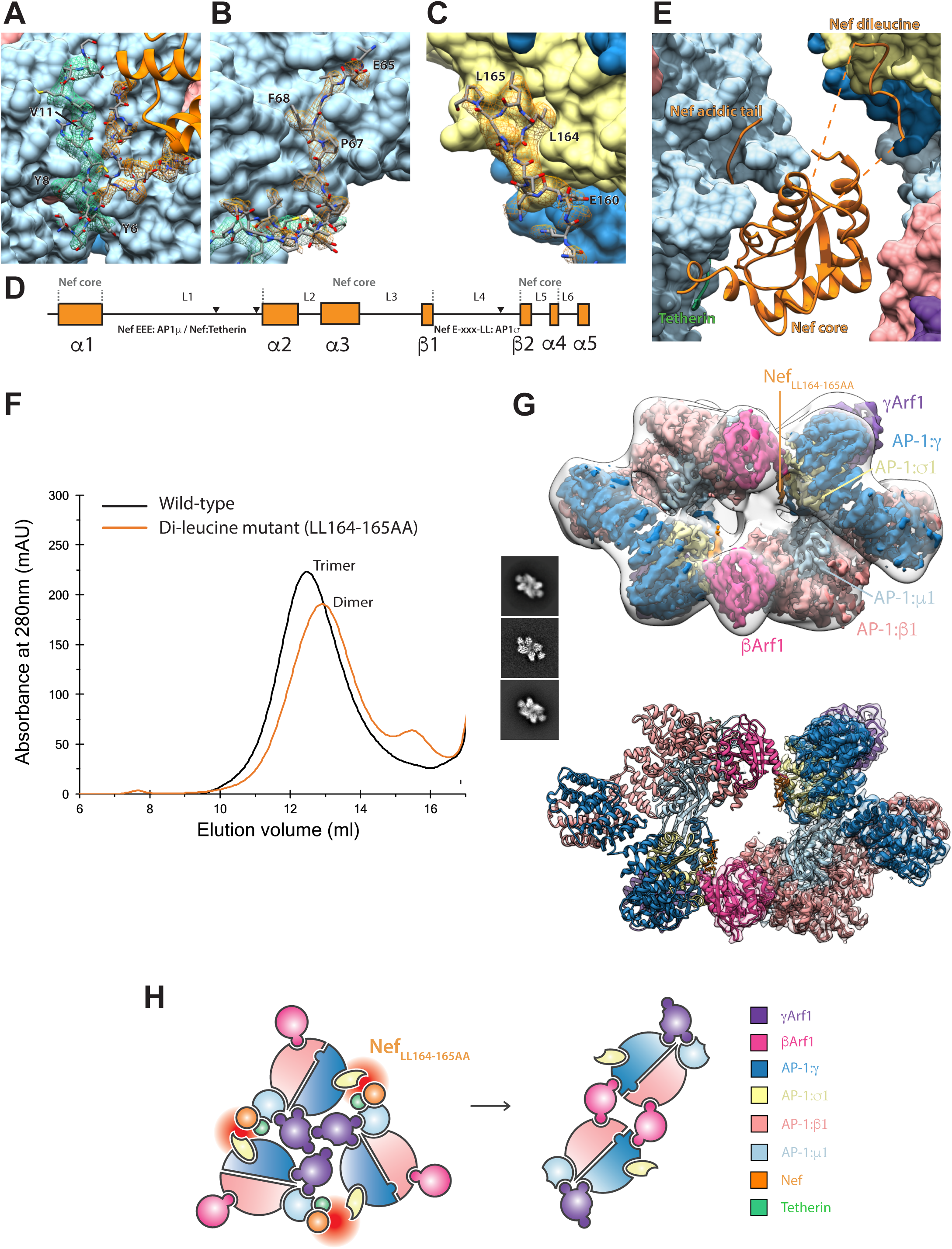
Nef bridging via the dileucine loop stabilizes the closed trimer. (A) An AP-1:μ1:Tetherin:Nef sandwich stabilises the Nef core to AP-1:μ1. (B) The canonical Nef acidic patch binds directly to AP-1:μ1. (C) The ExxxLL motif of Nef binds to AP-1:σ1. (D) These binding regions of Nef are found on the unstructured loops L1 and L4 but are stabilized in the trimeric assembly. (E) Consequently, the trimer architecture is closed by HIV-Nef at the AP1 trimeric dimer interface. (F) Knocking out the dileucine loop, tetherin-Nef^LL164-165AA^, responsible for binding AP-1:σ1 results in trimer destabilization and dimer formation as indicated by SEC. (G) The dileucine mutant adopts a dimeric oligomeric state as confirmed by cryo-EM. A 2D class and corresponding volume slice and projection are shown for reference. (H) The unstable trimer created by the knockout of the AP-1:σ1:LL interaction leads to a rearrangement into dimers driven by new βArf1:AP-1:γ interactions. See also Figure S1-3 and 4.

The γArf1, as well as the AP-1 core, are well-defined by the density (Figure S2B). The three γArf1 molecules contact one another through their β2-β3 turn, α4-β6 connection, and C-terminal helix (Figure 3A). The structure revealed multiple contacts between γArf1 α2 and α3 with the µ1-CTD (Figure S4A), confirming a previous report of a non-switch Arf1 region interacting with the related µ4 domain (Boehm et al., 2001). The three βArf1 molecules bound at the periphery of the complex are in an appropriate geometry to insert into the membrane via their N-terminal myristates, and the µ1-CTD binding site for MHC-I, tetherin, and other YxxΦ based signals is an unobstructed ∼20 Å from the membrane (Figure 1D). The γArf1 molecules are membrane-distal in this geometry, but Nef is poised to bind the membrane via its own myristoyl modification. Thus, at least nine points of contact between the closed assembly and membranes are evident, showing how this assembly could bind cargo at the TGN.

**Figure 3.**
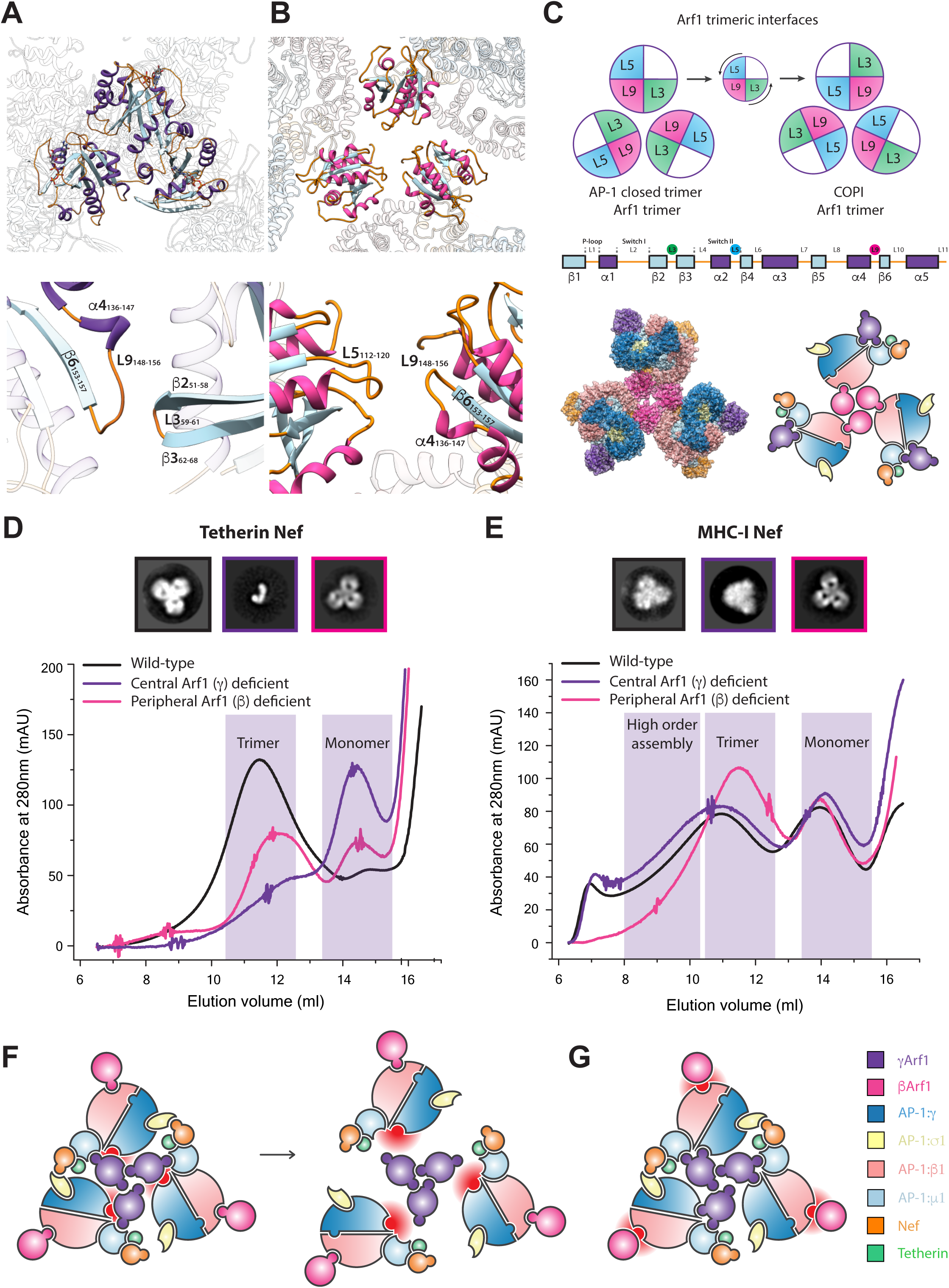
γArf1 and βArf1 control AP-1 oligomerization in a cargo dependent manner. (A) The binding loops of the closed AP-1 trimer are mediated through γArf1 loop 9 – loop 3 contacts. (B) Conversely the trimerisation architecture observed in COPI assemblies utilizes a rearrangement to loop 9 – loop 5 contacts. (C) The counterclockwise turn of the Arf1 assemblies allows these new contacts to be made, yet both still utilize loop 9. In the COPI-like Arf1 L9-5 trimeric architecture, assembly via βArf1 is sterically plausible. (D) In the presence of tetherin-Nef cargo Arf1 dependent trimerization of AP-1 is dependent on γArf1 but not βArf1. (E) In the presence of MHC-I-Nef cargo, AP-1 may trimerize via both γArf1 or βArf1. EM 2D class averages reveal an assembly in the presence of MHC-I-Nef cargo that thus may be assembled via βArf1 trimerization. (F) The knockout of the γArf1:AP-1:γ interface completely destabilizes tetherin trimers to monomer, but does not disrupt MHC-I assembly in the same manner. (G) The knockout of the βArf1:AP-1:β1 does not affect the tetherin trimer assembly but in the case of MHC-I trimers allows only assembly via the γArf1 trimer. Thus, MHC-I trimers can assemble via both γArf1 and βArf1 oligomerization. See also Figure S4.

### Nef bridging uniquely stabilizes the closed trimer

The structure described above suggested that the strength of the µ1-CTD-LL site bridge might control the stability of the closed trimer. The properties of the tetherin-Nef^LL164-165AA^ assembly were therefore investigated. AP-1:Arf1:Tetherin-Nef^LLAA^ migrated on SEC as an apparent dimer (Figure 2F), confirming that the Nef dileucine sequence is essential for closed trimer formation. The structure of the AP-1:Arf1:Tetherin-Nef^LLAA^ dimer was determined at 4.27 Å resolution by cryo-EM (Figure 2G, Figure S1B, 2C, 3G-I). AP-1 is in the hyper-unlocked conformation, as seen in the closed trimer, thus the change in oligomerization is not driven by a change in AP-1 monomer conformation. As seen previously in the crystal structure of the unlocked AP-1:Arf1 dimer determined in the absence of Nef or cargo (Ren et al., 2013), two βArf1 molecules cross-link the dimer, while the AP-1 complexes themselves do not contact one another. However, the hyper-unlocked conformation of AP-1:Arf1:Tetherin-Nef^LLAA^ cross-links through a unique βArf1-AP-1:γ contact that is not seen in the crystal structure of the unlocked conformation (Figure S4D). This new contact allows for dimerization of the hyper-unlocked conformation as the γArf1-AP-1:γ interaction present in the unlocked conformation is no longer available for crosslinking. Residues ∼160-166 of Nef^LLAA^ were observed bound to σ1-γ despite the absence of the Leu side chains (Figure S4F). Thus, Nef^LLAA^ is still capable of binding to AP-1 at the high concentrations used for reconstitution, but the interaction is too weak to tether two AP-1 subunits in the closed trimer assembly. These data confirm that the closed trimer assembly is induced by Nef dileucine-mediated bridges.

### Alternative Arf1 trimers lead to alternative assemblies

We previously reported that AP-1:Arf1:Nef complexes could form open trimeric assemblies that strongly promoted clathrin basket formation (Shen et al., 2015). The open trimers are only connected by Arf1 trimerization at the center, and the ensuing dynamics and heterogeneity of these structures has limited attainable resolution to a 17 Å reconstruction that is presumably a superposition of multiple conformations. Recently, the cryo-electron tomography (cryo-ET) and sub-volume averaged 9 Å structure of an Arf1 trimer bound to the related coat complex COPI was reported (Dodonova et al., 2017). The AP-1-γArf1 closed trimer described above is distinct from the COPI-Arf1 trimer, although they share the use of the "L9" (α4-β6) loop (Figure 3A-B). We modeled combinations of βArf1 and γArf1 bound to hyper-unlocked AP-1 trimerized via either the closed trimer or the COPI Arf1 trimer geometry. AP-1 bound to βArf1 in the COPI geometry was sterically possible and competent to bind membranes in the context of known AP conformations with hyper-unlocked AP-1 subunits (Figure 3C). This suggested to us that the closed trimer was driven by γArf1, and the open trimer by βArf1.

We disrupted the two Arf1 interfaces on AP-1 using the previously described mutants AP-1^ΔβArf1^ and AP-1^ΔγArf1^ (Ren et al., 2013). In the context of tetherin-Nef, which forms predominantly closed trimer assemblies under the conditions tested, AP-1^ΔβArf1^ migrated almost normally on SEC, while AP-1^ΔγArf1^ shifted sharply towards a monomeric distribution (Figure 3D), as predicted by the closed trimer structure. In contrast, in the MHC-I-Nef sample, which is known to consist of open trimer and higher order structures (Shen et al., 2015), AP-1^ΔγArf1^ behaved identically to wild-type (Figure 3E). In contrast, AP-1^ΔβArf1^ bound to MHC-I-Nef (Figure 3E) migrated in the position expected for the closed trimer (Figure 3D). Thus, the AP-1^ΔβArf1^ mutation switches what would have been open MHC-I trimers into closed trimers. These data strongly support the concept that alternative Arf1 trimerization and the use of alternative Arf1 contact sites on AP-1 controls whether the assembly formed is open or closed.

### Nef polyproline strand dynamics mediates cargo selectivity

The Nef core is inherently mobile in the closed trimer EM density (Figure S9), yet its presence is critical for bridging and stabilizing the trimer. The observation suggested to us that Nef flexibility might positively regulate formation of the closed trimer. We used HDX-MS (Figure S6) to compare the dynamics of Nef in the context of complexes of the tetherin-Nef and MHC-I-Nef constructs with µ1-CTD. Consistent with the crystal structure of the MHC-I-Nef: µ1-CTD complex (Jia et al., 2012), which showed extensive interactions between the polyproline region and MHC-I, peptides overlapping with this segment (residues 66-80; Figure 4A, B) were well-protected (< 20 % exchange at 10 s; Figure S6B). This region was less protected in the tetherin complex (∼50 % exchange at 10 s; Figure S6A). Elsewhere in Nef, there were relatively few differences in protection. These data show that Nef is held rigidly to the µ1-CTD in the MHC-I complex, while it is bound more flexibly in the tetherin complex. This flexibility could facilitate formation of the asymmetrically bridged closed trimer (Figure 1C, 4C) by lowering the energetic cost for Nef to bridge from µ1-CTD in one AP-1 complex to the σ subunit of the adjoining AP-1 complex.

**Figure 4.**
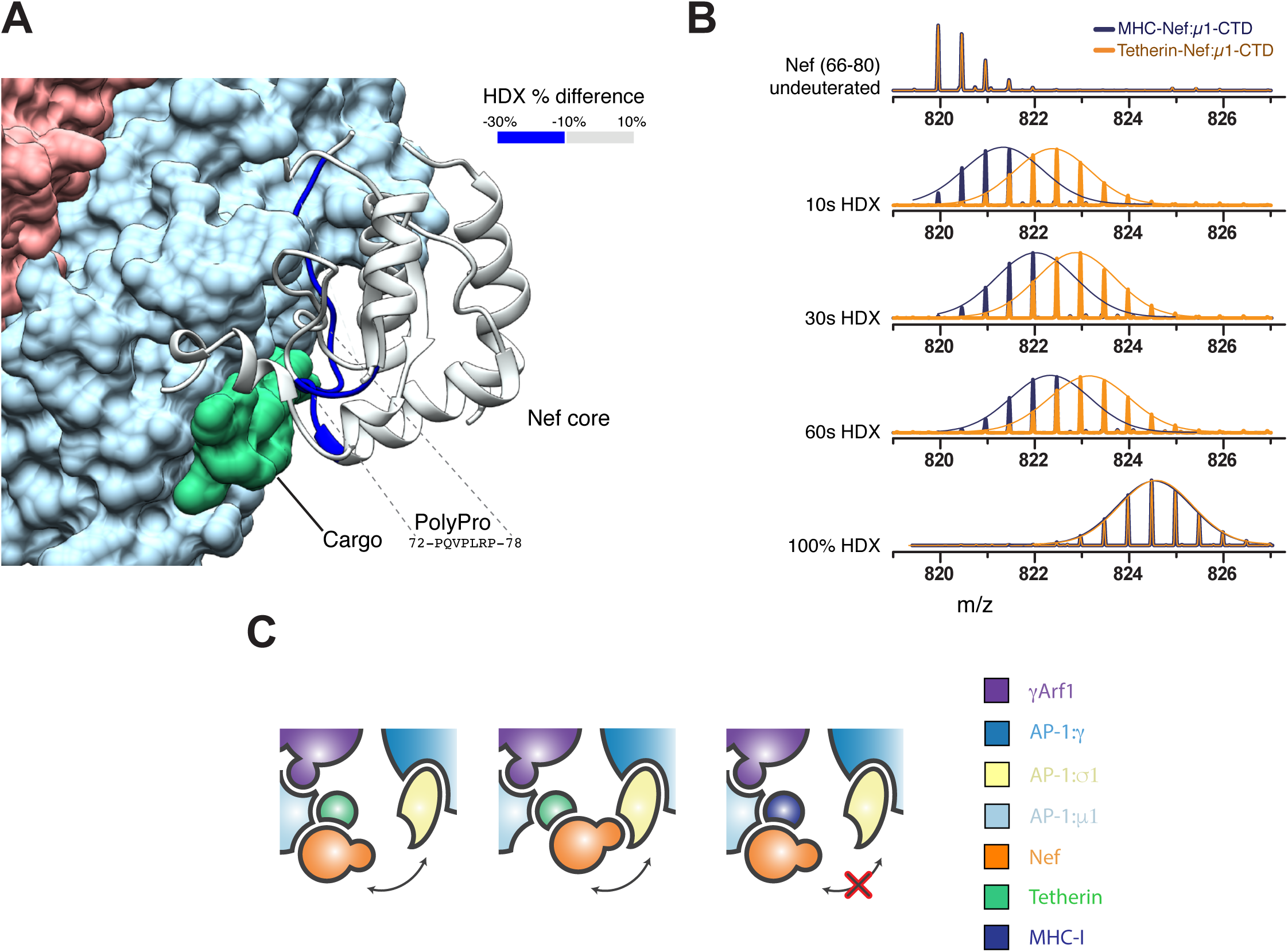
Cargo modulates Nef dynamics. (A) Mapping of the changes in deuteration levels of Nef between tetherin-Nef:AP1:μ1-CTD and MHC-Nef:AP-1:μ1-CTD. The HDX-MS Difference at 10s is overlaid onto the cryoEM structure of the AP-1:Arf1:tetherin-Nef trimer. (B) Mass spectra of the peptide from Nef (66-80) in tetherin-Nef (orange) or MHC-Nef (blue) upon AP-1:μ1-CTD binding. A Gaussian fit is used to represent the distribution of peak heights of the ion peaks across the m/z values, and overlay with mass spectra (straight line). (C) The stronger binding of Nef to AP-1:μ1 in the presence of MHC-I cargo may prevent Nef from reaching its AP-1:σ1 binding site. See also Figure S5.

### Phosphoregulation of the closed trimer

We sought to determine whether the closed trimer is functionally important in tetherin downregulation. HIV-1 NL4-3 Nef is considered functionally inert in tetherin counteraction, while HIV-1 group O Nefs are quite effective (Kluge et al., 2014). This functional difference has been mapped to a handful of residues surrounding the dileucine signal (Figure 5A). We used GST pull-down assays to determine the ability of NL4-3 and O-MRCA Nef proteins to interact with a µCTD-deleted form of the AP-1 core (AP-1 core^ΔµCTD^), which is capable of interacting with cargo via the dileucine motif but not the YxxΦ motif (Jia et al., 2014). NL4-3 Nef bound to the truncated construct (Figure 5B), consistent with the observation of LL binding to the σ1-γ in the cryo-EM structure of the closed trimer. O-MRCA Nef bound robustly to AP-1 core^ΔµCTD^ as well (Figure 5B). This is potentially consistent with a functional role for an O-Nef:AP-1 interaction in tetherin downregulation, but begs the question why a greater difference between NL4-3 and O-Nefs was not observed. Given the presence of a Ser adjacent to the LL motif in NL4-3 and most other M-Nefs, but not O-Nefs (Figure 5A), we considered whether this residue might be subject to phosphoregulation. Ser169 is predicted to contain a consensus phosphorylation site for casein kinase I (Hornbeck et al., 2015). We found that CKI phosphorylated Ser169 uniquely and stoichiometrically *in vitro* (Figure 5C). CKI phosphorylated NL4-3 Nef does not bind to AP-1 core^ΔµCTD^ (Figure 5A-B). Consistent with the dependence of closed trimer formation on LL engagement, phospho-Nef does not manifest a closed trimer peak on SEC, but rather redistributes to aggregate, open trimer, and monomer peaks (Figure 5D). These data show that closed trimer formation by NL4-3 Nef is subject to phosphoinhibition via a site which is absent in O-Nefs.

**Figure 5.**
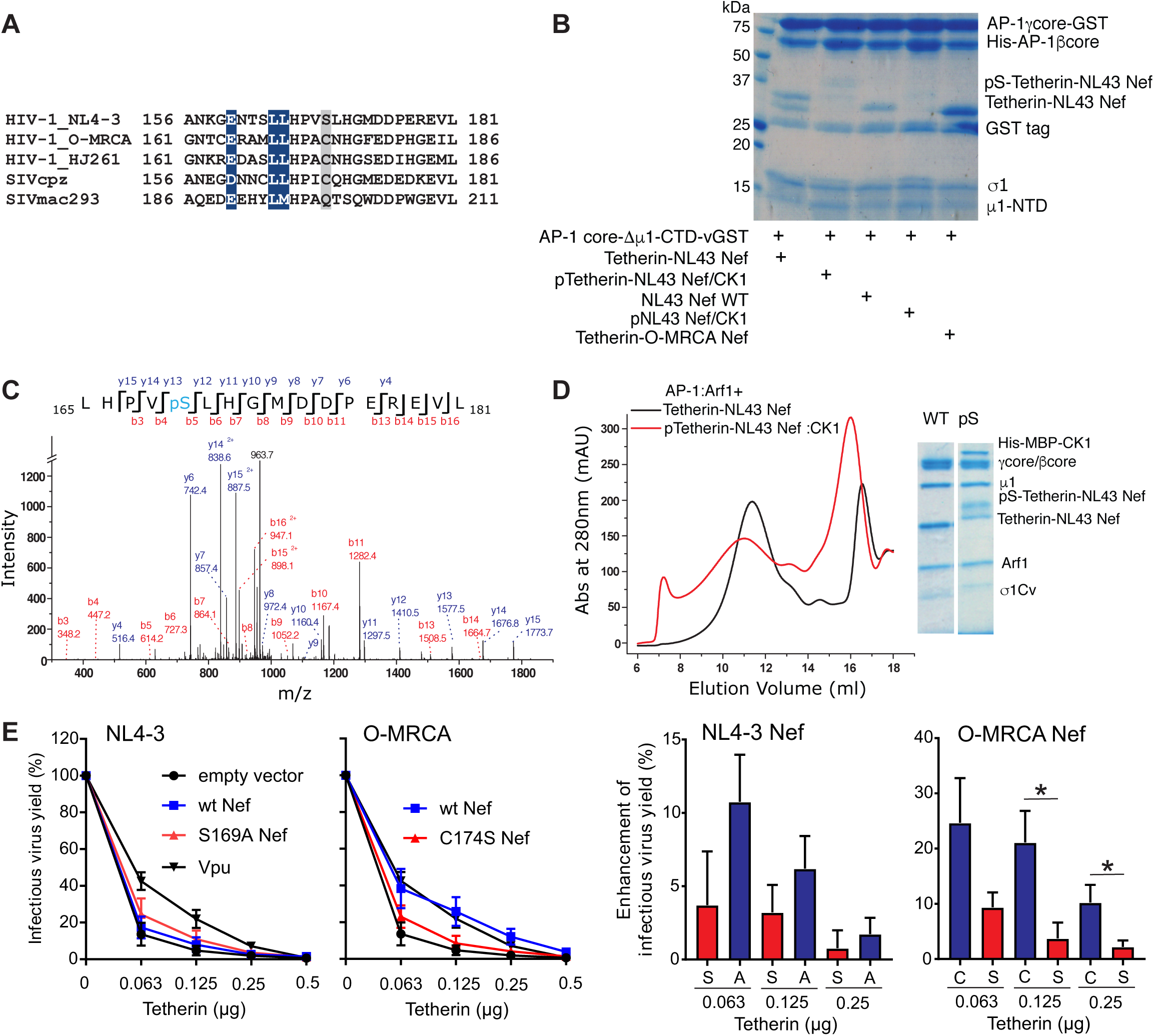
Phosphorylation of NL4-3 Nef by CK1 disrupts the interaction between AP-1 σ1 and Nef dileucine motif. (A) Amino acid alignment of Nef proteins in the D/ExxxLL motif region. The dileucine motifs that are conserved in all Nef strains are colored in blue. The residues aligned with NL4-3 Nef Ser169 predicted CK1 phosphorylation site (Hornbeck et al., 2015) are highlighted in gray. (B) Collision-induced dissociation (CID) fragmentation spectrum of the phosphorylated peptide in tetherin-Nef treated by CK1. Ser169 in NL4-3 Nef is phosphorylated. (C) GST pull down result suggest that Nef phosphorylation by CK1 largely reduces Nef dileucine motif binding to AP-1. AP-1 core^μCTD^-GST is selected to represent the interaction with the Nef dileucine motif. The phosphorylation of Nef largely reduces the binding to AP-1core^μCTD^-GST on the beads. (D) Gel filtration and SDS gel analysis showing the contents of trimer peak. The phosphorylation of Nef disrupts the closed trimer assembly. (E) The effect of Nef differing in the Ser phosphorylation site on infectious virus release in the presence of tetherin. Infectious virus yield from 293T cells co-transfected with an HIV-1 NL4-3 ΔVpuΔNef construct and vectors expressing the indicated *nef* alleles or human tetherin was determined by infection of TZM-bl cells (right). Data show mean percentages (±SEM) relative to those detected in the absence of tetherin (100%) obtained in five independent experiments. Results obtained for NL4-3 Vpu are shown for comparison. The bar diagrams show the increase in virus production in the presence of Nef compared to the vector control. Stars refer to the difference from the EGFP control panel. ^*^p < 0.05, ^**^p < 0.01.

To directly determine the relevance of this Ser phosphorylation site for the anti-tetherin activity of Nef, we introduced mutations of S169A and *vice versa* C174S in the NL4-3 and O-MRCA Nef proteins, respectively. The parental NL4-3 Nef only marginally increased infectious virus yield from 293T cells after cotransfection of a ΔVpuΔNef HIV-1 NL4-3 proviral vector with fixed quantities of Nef expression construct and different doses of tetherin expression plasmid. Mutation of S169A increased its capability to counteract tetherin, although it remained substantially less effective than Vpu or the O-MRCA Nef (Figure 5E). This is in agreement with previous data showing that S169C only conferred full anti-tetherin activity to the HIV-1 JC16 Nef in combination with a S163I change in the third variable residue of the ExxxLL motif (Gotz et al., 2012). Mutation of C174S, however, almost fully disrupted the anti-tetherin activity of O-MRCA Nef (Figure 5E) supporting a role of phosphoregulation in the ability of Nef to counteract tetherin.

## Discussion

Nef is an important determinant of HIV infectivity and pathogenesis, and exerts many, if not most, of its effects by hijacking the coated vesicle trafficking pathways of infected cells. We previously showed that NL4-3 Nef hijacks AP-1 in a startlingly complex manner (Shen et al., 2015). Here, we build on this work to show how different AP-1 hijacking mechanisms are used by cargoes undergoing different fates, and by Nefs of the M and O groups of HIV-1 groups that are known to counteract the host restriction factor tetherin by divergent means. Our previous structural study revealed that tetherin and NL4-3 Nef stabilize a closed AP-1 trimer, while MHC-I and NL4-3 Nef promote an open conformation. Both structures are capable of docking onto membranes but only the open trimer has a geometry matching that of the clathrin lattice (Shen et al., 2015). The role of the closed trimer was mysterious, with progress limited by the intermediate resolution of the previous study. Here, we obtained a large data set on a Titan Krios and utilized a signal subtraction and intra-trimer subunit averaging strategy accounting for local symmetry deviations to obtain an atomistic reconstruction of the closed trimer.

The atomistic structure of the closed trimer, and comparison to related structures, clarified two points central to functional analysis. First, a large section of the Nef LL loop was visualized in the density. This loop could be uniquely assigned to a Nef molecule bound *in trans* to the µ1-CTD of a different AP-1 complex within the trimer. This established that Nef binds *in trans* between the LL binding site and the µ1-CTD, and suggested that the ability of Nef to bridge these two sites *in trans* could be the driving force for closed trimerization. Second, the improved resolution, together with the publication of a higher resolution structure of an Arf1-mediated trimeric interface in the related COPI coat, suggested that the open and closed trimers use Arf1 contacts differently. These higher resolution insights made it possible to functionally dissect the determinants for closed trimerization.

The LL-µ1-CTD bridge model predicted that the closed trimer must be held together by strong contacts on both the LL and µ1-CTD sides, with sufficient affinity to drive formation of an oligomer that would otherwise be energetically unfavorable. We thus expected that mutation of the LL sequence to dialanine (AA) would block formation of the closed trimer. Confirming this idea, we found that tetherin and Nef^LLAA^ forms a mixture of monomers and dimers, but not closed trimers. We went on to determine an near-atomistic cryo-EM structure of the tetherin-Nef^LLAA^ dimer. The LLAA loop density could be visualized in this structure, showing that the residual affinity of the LLAA mutant loop was adequate to bind at the high concentrations used to generate the sample, but inadequate to drive closed trimerization. Subsequently, we found that CK1 phosphorylation of Ser169, which immediately follows the LL sequence in NL4-3 Nef, also weakens binding to the LL site. In this case, a mixture of aggregates, open trimers, and monomers replaces closed trimers. While it is still unclear what controls the population of dimers and open trimers when closed trimerization is blocked, these data firmly establish LL-µ1-CTD bridging as the mechanism for stabilizing closed trimers.

These and previous (Shen et al., 2015) data and the recent 9 Å cryo-ET structure of the COPI coat (Dodonova et al., 2017) highlight the conserved and ancient role of Arf1 as a building block and integral component of vesicular coats. The conformations of the open and closed AP-1 trimers are so different that it seems impossible to rationalize them in terms of a single mode of Arf1 trimerization. Differential trimerization was confirmed using a set of mutations in the Arf1 binding site of the β1 and γ subunits of AP-1 (Ren et al., 2013). Blocking the γ site converts the tetherin complex from a closed trimer to a monomer, while the while β1 site mutant had little effect. Disruption of the γ site has little effect on the MHC-I complex, but strikingly, blocking the β1 site converts it from open trimer to closed. Previously, we showed how the presence of multiple Arf1 binding sites on AP-1 could result in the cross-linking of AP-1 into larger polygons (Shen et al., 2015). We now find that the multiplicity of binding sites leads to the ability to trimerize in ways that lead to sharply different functions.

SIV/HIV is believed to have made multiple independent crossings of the simian/human species barrier. SIV downregulates tetherin via Nef hijacking of AP-2, a mechanism that depends on the presence of the sequence (G/D)DIWK in simian tetherins (Zhang et al., 2009). The inability of SIV Nefs to downregulate human tetherin represents one of the main barriers to interspecies transmission (Neil et al., 2008; Van Damme et al., 2008). The pandemic M-group HIV-1 uses Vpu to downregulate tetherin (Neil et al., 2008; Van Damme et al., 2008), and this effect appears to occur through retention of Vpu and tetherin in the TGN (Dube et al., 2009; Jia et al., 2014). However, O-group viruses and a few others appear to have evolved a distinct Nef-dependent but (G/D)DIWK-independent mechanism, which also entails retention in the TGN (Kluge et al., 2014). The closed trimer structure, which is competent to bind membranes but we expect not to drive clathrin assembly, suggests a structural explanation for TGN retention (Figure 6). In this model, the closed trimer binds to tetherin, Nef, and Arf1 at the TGN membrane, but does not progress efficiently to clathrin coat formation because the geometry of the closed trimer does not match that of the clathrin lattice. One apparent flaw of this model is that we observed closed trimer formation with both NL4-3 and O-MRCA Nefs, despite that NL4-3 Nef is known to be ineffective in driving TGN retention of tetherin. We noticed NL4-3 and most other M-group Nefs, but not O-Nefs, are predicted to be phosphorylated on the LL loop by CK1, and confirmed the phosphorylation of NL4-3 Nef experimentally. Notably, the Ser phosphorylation site is found in most HIV-1 M Nefs but generally absent in Nef proteins of SIVcpz, the direct precursor of pandemic HIV-1 strains. In agreement with the present results, it has been shown that substitution of S169C is critical for effective tetherin antagonism by HIV-1 Nefs but does not significantly affect other Nef functions (Gotz et al., 2012; Kluge et al., 2014). Our results thus explain how M-Nefs were functionally inactivated with respect to TGN retention of tetherin because HIV-1 M strains evolved Vpu as potent tetherin antagonist during adaption to the new human host.

**Figure 6.**
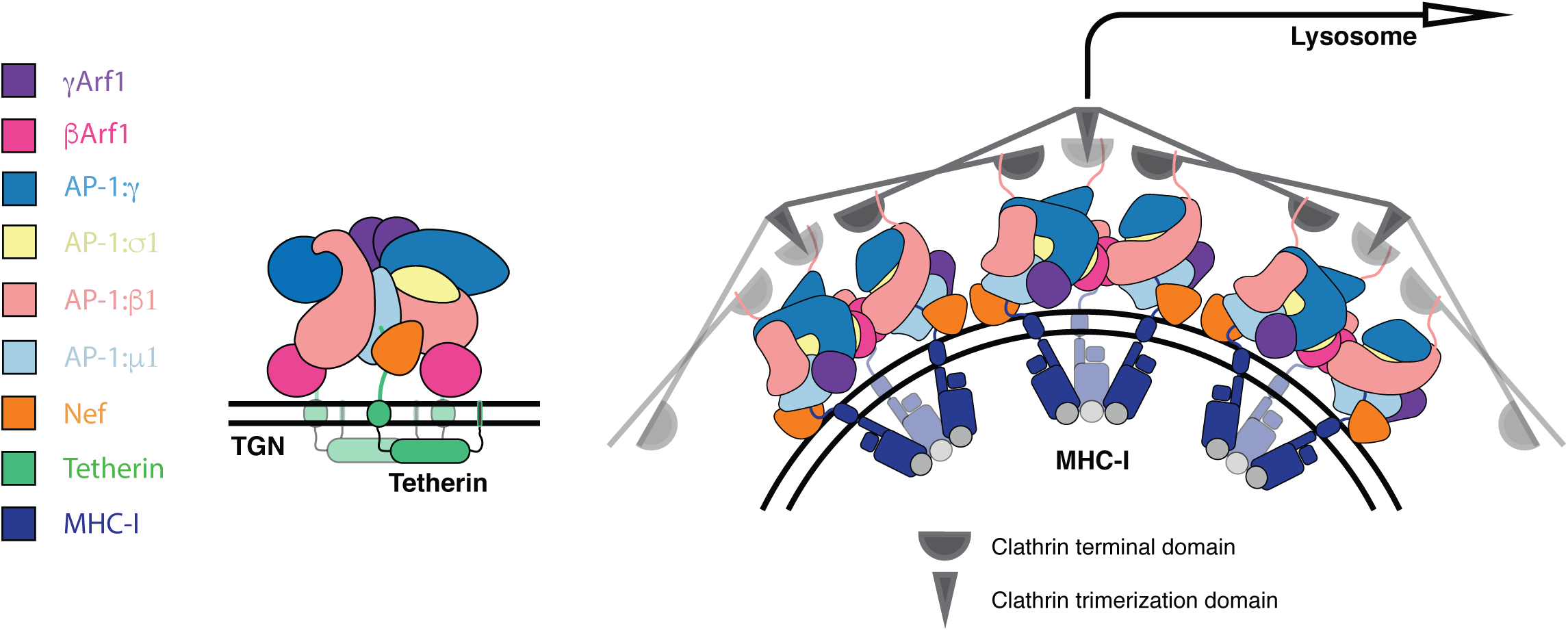
Model for allosteric regulation of Nef target fate. Tetherin cargo loaded AP-1:Arf1 trimers are retained at the trans golgi network in a trimeric architecture not competent to assemble into higher species and thus recruit clathrin. MHC-I cargo loaded AP-1:Arf1 trimers may assemble through allosterically induced architectural and assembly changes that permits higher order assembly, clathrin recruitment and trafficking to the lysosome. See also Figure S6.

Considerable effort has gone into characterizing how the allosteric unlocking of single AP monomers regulates their ability to bind to cargo (Jackson et al., 2010; Kelly et al., 2008), membranes (Collins et al., 2002; Ren et al., 2013), and clathrin (Kelly et al., 2014). We have now shown that in the context of HIV-1 infection, allosteric regulation also occurs at the level of alternative trimerization of AP-1 complexes. It remains to be clarified why TGN retention, as opposed to lysosomal degradation, seems to be advantageous for HIV-1. Possibly, retention in a subdomain of the TGN might help prevent the tetherin sorting pathway from intersecting that of its substrate, HIV-1 Env. Beyond the implications for HIV-1 Nef, the larger implication of these results is that higher order assemblies of vesicle coat adaptors are remarkably plastic, and that this plasticity can be harnessed for regulatory purposes.

### Experimental Procedures

#### Plasmid construction

The His_6_- and GST-tagged AP-1 constructs were previously described (Shen et al., 2015). MHC-I (338-365)-NL4-3 Nef (no linker) (Jia et al., 2012), Tetherin (1-21)-10aa linker-NL4-3 Nef and Tetherin (1-21) -10aa linker-O-MRCA (Kluge et al., 2014) Nef were expressed as TEV-cleavable N-terminal His_6_ fusions. The rat CK1δ (1-317) gene was codon-optimized, synthesized using gblocks (Integrated DNA Technologies, Coralville, Iowa), and then subcloned into LIC 2C-T vector, which was expressed as TEV-cleavable N-terminal His_6_–MBP fusion. Bicistronic CMV promoter-based pCG expression vectors coexpressing parental or mutant Nef proteins were generated as described (PMID: 25525794). All constructs used in this study are listed in Table S1.

#### Protein purification

The AP-1 complexes were expressed in BL21 (DE3) pLysS (Promega, Madison, WI) strains and induced with 0.3 mM isopropyl-β-D-thiogalactopyranoside (IPTG) at 20^°^C overnight. The cells were lysed by sonication in 50 mM Tris at pH 8.0, 300 mM NaCl, 10% glycerol, 3 mM β-mercaptoethanol (β-ME), and 0.5 mM phenylmethanesulfonyl fluoride (PMSF). The clarified lysate was first purified on a Ni–nitrilotriacetic acid (NTA) column (Qiagen, Valencia, CA). The eluate was further purified on glutathione– Sepharose 4B resin (GE Healthcare, Piscataway, NJ). After TEV cleavage at 4^°^C overnight, the sample was concentrated and then loaded onto a HiLoad 16/60 Superdex 200 column (GE Healthcare) in 20 mM Tris at pH 8.0, 200 mM NaCl, and 0.3 mM tris(2-carboxyethyl)phosphine (TCEP). The sample fractions were pooled together, adjusted to 30 mM imidazole, and passed through 1 ml of glutathione–Sepharose 4B and then onto a Ni-NTA column (Qiagen) to capture the residual GST-and His-tag fragments. The sample was adjusted to 20 mM Tris at pH 8.0, 200 mM NaCl, and 0.3 mM TCEP by buffer exchange in the concentrator.

His_6_-tagged tetherin-Nef and His_6_-tagged Arf1 constructs were expressed in BL21 (DE3) Star cells and induced with 0.3 mM IPTG at 20^°^C overnight. The cell pellet was lysed by sonication and purified on a Ni-NTA column in 50 mM Tris at pH 8.0, 300 mM NaCl, 20 mM imidazole, 5 mM MgCl_2_, 3 mM β-ME, and 0.5 mM phenylmethanesulfonyl fluoride (PMSF). The proteins were eluted with 300 mM imidazole and then loaded onto a HiLoad 16/60 Superdex 75 column (GE Healthcare) in 20 mM Tris at pH 8.0, 200 mM NaCl, 5 mM MgCl and 0.3 mM TCEP. The sample fractions were pooled and proteins were quantified by molar absorption measurements.

#### AP-1:Arf1:tetherin-Nef complex assembly

The recombinant AP-1 core was mixed with Arf1 and tetherin-Nef proteins at a molar ratio of 1:4:6 and then incubated with 1 mM GTP at 4^°^C overnight. The mixture was then subjected to a Superose 6 10/100GL column in 20 mM Tris at pH 8.0, 200 mM NaCl, 5 mM MgCl_2_, and 0.5 mM TCEP. For the purposes of EM analysis, fractions corresponding to AP-1:Arf1:tetherin-Nef complexes were pooled and concentrated for cryoEM analysis.

#### Cryo-electron microscopy

Samples for data collection were prepared on 1.2/1.3 C-flat holey carbon copper grids that were prepared by plasma cleaning for 10 s using a Solarus plasma cleaner (Gatan Inc., Pleasanton, CA). Reconstituted complex at 0.7 mgmL^-1^ was applied in a 3 µL drop to one side of the grid and plunge frozen into liquid ethane using a Vitrobot mark VI. The humidity was controlled at 100%, 22 ^°^C and the grids blotted with Whatman #1 paper using a blot force of 8 for between 3.5 and 4.0 seconds. 2,200 micrographs were collected on a Titan Krios (FEI; BACEM UCB) at a nominal magnification of 22,500X and a magnified pixel size of 1.067 Å pix^-1^. The dose rate was 6.57 e^-^/Å^2^/sec at the sample and achieved with an illuminated area sufficient to cover one hole and the surrounding carbon support. Data collection was carried out using SerialEM with one exposure per hole and focusing for each exposure, the defocus range was set to collect between 0.75 and 2.00 µm defocus. Movies were acquired on a K2 direct electron detector (Gatan) in super-resolution counting mode with a dose fractionated frame rate of 250 ms and total collection time of 9500 ms.

AP1:Arf1:tetherin-Nef (L2A) cryo grids were prepared identically to the WT AP1:Arf1:tetherin-Nef sample. 1,989 micrographs were collected on a Titan Krios at a nominal magnification of 22,500X and a magnified pixel size of 1.067 Åpix^-1^. The dose rate was 7.09 e^-^/Å^2^/sec at the sample. Data collection was carried out using SerialEM as reported for the WT AP1:Arf1:tetherin-Nef sample except with a defocus range between 0.75 and 2.50 µm and a dose fractionated frame rate of 200 ms and total collection time of 5600 ms. Full details of data sets are listed in Table S2.

#### Image processing

Movie data were processed using MotionCor2-1.0.0 (Li et al., 2013) with internal 2-fold Fourier binning, discarding the initial 2 frames, 5 by 5 patch based alignment and dose weighting up to the total exposure of 62.4 e^-^/Å^2^. The contrast transfer function of full dose non-weighted micrographs was estimated using Gctf-v1.06 (Zhang, 2016). 252,212 particles were picked from non-dose-weighted micrographs using Gautomatch-v0.50, particles were extracted in a 384 pixel box and subjected to three rounds of 2D classification at 3-fold binning in Relion-2.0 (Kimanius et al., 2016) resulting in 64 classes with 209,816 selected particles. Ab-initio 3D classification was performed in cryoSPARC (Punjani et al., 2017) extracting a monomer structure with 75,442 particles that could be refined to 4.23 Å and over three 3D classifications and subset selections leading to a closed trimer structure with 61,948 particles reaching 6.25 Å. The closed trimer particle angles and reference volume were refined in Relion-2.0 to 4.27 Å in C1. To account for the pseudo-symmetry present in the complex due to local monomeric subunit movement we used Relion localized reconstruction (localrec) (Ilca et al., 2015) through in house written scripts for extracting monomeric subunits of the closed trimer as new subparticles (Figure S2). The trimer subunits (containing the AP1 core, two copies of Arf1 and two copies of Nef) were localized, recentered and extracted in new 224 pixel boxes. For each subunit extraction, a vector describing the center of the trimeric refinement volume to the center of mass of the AP1 subunit was defined in UCSF Chimera (Pettersen et al., 2004). Coordinates of the subunit were rigid-body fitted to the vector-localized subparticle and used to segment the map using *scolor* by selection within 25 Å of Cα atoms. The remaining density was softened by multiplication with at 3 σ mask extended and softened by 4 and 9 pixels, this was used for signal subtraction to leave only the subunit of interest. Using the Relion refinement parameters and vectors, localrec was used to localize the subparticles in every whole particle image with and without signal subtraction. These were extracted as new particles, with recentred origins. The subparticle extraction was repeated for each monomer of the closed trimer structure resulting in 185,787 extracted subparticles. Two rounds of 2D classification cleaned the subparticle image stack to 139,172 particles. The subparticles were refined in Relion-2.1 (beta-1) to 3.93 Å with a measured map B-factor of -164 Å^2^. A masked 3D classification into 3 classes with Tau 10 and no angular searches produced a particle stack of 53,841 subparticles that were refined to 3.73 Å with a measured map B-factor of -137 Å^2^. For focused 3D classification of the Nef density a mask was created around the segmented density of Nef and 3D classification performed on the density inside this mask without sampling of the angles obtained from the 3D refinement, into 3 classes with Tau 16. This successfully characterized the high, partial or low occupancy of the Nef density.

For the AP1:Arf1:tetherin-Nef (L2A) sample, movie data were processed using MotionCor2-1.0.0 and the contrast transfer function of full dose non-weighted micrographs was estimated using Gctf-v1.06. 162,309 particles were picked from non-dose-weighted micrographs using Gautomatch-v0.50. Particles were extracted in a 384 pixel box and subjected to a round of 2D classification at 4-fold binning in Relion-2.1 (beta-1). Seventeen selected classes were then used for template based Auto-picking in Relion-2.1 (beta-1), which yielded 399,032 particles. These particles were subjected to 4 iterative rounds of 2D classification in Relion-2.0, resulting in 51 classes with 162,509 selected particles. Ab-initio 3D classification was performed in cryoSPARC, extracting two dimer structures with 17,994 particles and 42,902 particles that could be refined to 8.64 Å and 7.80 Å, respectively. The two dimer particle angles and reference volumes were further refined in Relion-2.0 to 7.31 Å and 6.72 Å, respectively, in C1. Similar to the wild-type AP1:Arf1:tetherin-Nef, the L2A dimer structure presented pseudo symmetry due to local subunit movement. Localized reconstruction subparticle extraction was performed in Relion on the of the 6.72 Å dimer subunit as described above. Subparticles containing the AP1 core and two copies of Arf1 were localized, centered and extracted in new 192 pixel boxes as shown in Figure S2. 85,804 subparticles were subjected to a round of 2D classification leading to 85,529 particles. The subparticles were refined in Relion-2.1 (beta-1) to 4.27 Å with a measured map b-factor of -151 Å^2^.

Map validation was performed by estimating local resolution with Relion. The particle orientation distribution scores were calculated using cryoEF (Naydenova and Russo, 2017).

#### Modeling

Modeling for the closed trimer monomeric subunit was performed in UCSF Chimera (Pettersen et al., 2004) using starting models from PDB entries 4P6Z, 4HMY and 4EN2, followed by manual adjustment in Coot (Emsley et al., 2010). Structure refinement using Phenix real space refinement (Adams et al., 2010) was used to improve the fit to the density and model geometry. Map-to-model comparison in phenix.mtriage validated that no overfitting was present in the structures and emRinger scores were calculated (Barad et al., 2015) to identify regions requiring remodeling in Coot (Emsley et al., 2010) before a final 5 cycle real space refinement in Phenix. Atomic displacement factor refinement was used to calculate the residue B-factors. The final model map cross correlation was 0.75, with a map-vs-model FSC_0.5_ of 3.87 Å and an average model B-factor of 26.5 Å^2^ for the AP1:Arf1:tetherin-Nef trimer. The final model map cross correlation was 0.77, with a FSC_0.5_ of 4.33 and an average B-factor of 83.2 Å^2^ for the WT AP1:Arf1:tetherin-Nef (L2A) dimer. Model geometry was validated using MolProbity (Davis et al., 2007). All map and model statistics are detailed in Table. S2.

#### HDX-MS

After the AP-1 trimer was separated on a Superpose 6 column, the trimer fraction was pooled and concentrated to 20 µM. Amide HDX-MS was initiated by a 20-fold dilution of AP-1 trimer complex into a D_2_O buffer containing 20 mM HEPES (pD 7.2), 200 mM NaCl, 2 mM MgCl_2_ and 0.2 mM TCEP at room temperature. After intervals of 10 s-60 s, exchange was quenched at 0 ^°^C with the addition of ice-cold quench buffer (400 mM KH_2_PO_4_/H_3_PO_4_, pH 2.2). The samples were then injected onto an HPLC system (Agilent 1100) with in-line peptic digestion and desalting. Desalted peptides were eluted and directly analyzed by an Orbitrap Discovery mass spectrometer (Thermo Scientific). Peptide identification was carried out by running tandem MS/MS experiments. A Proteome Discoverer 2.1 (Thermo Scientific) search was used for peptide identification. Mass analysis of the peptide centroids was performed using HDExaminer (Sierra Analytics, Modesto, CA), followed by manual verification of each peptide. The deuteron content was adjusted for deuteron gain/loss during digestion and HPLC. Both nondeuterated and fully deuterated complexes were analyzed. Fully deuterated samples were prepared by three cycles of drying and re-solubilization in D_2_O and 6 M guanidinium hydrochloride.

#### Nef phosphorylation by CK1

40 µM Tetherin-NL4-3 Nef or NL4-3 Nef was incubated with His-MBP-CK1 (6 µM) in the sample buffer consisting of 4 mM MgCl_2_, 1 mM ATP at 30 °C for 2 hr. 5 µl of 20 µM sample was diluted into 195 µl of 0.05% TFA, then injected into Agilent 1100 HPLC with in-line peptic digestion. Digested peptides were identified and characterized by tandem mass spectrometry (MS/MS) in an Orbitrap Discovery mass spectrometer. Proteome Discoverer 2.1 software was used to search the phosphorylation site in NL4-3 Nef.

#### GST pull-down assay

20 µg of recombinant AP-1 core^ΔµCTD^-GST was immobilized on 30 µl glutathione Sepharose and incubated with the wild type or phosphorylated Nef samples at 4 °C overnight in 20 mM Tris pH 8, 200 mM NaCl, 0.1 mM TCEP. The beads were washed 4 times, mixed with 40 µl of SDS buffer and boiled for 3 min. 15 µl of each sample was subjected to reducing SDS/PAGE.

#### Tetherin antagonism

To determine the capability of Nef to antagonize tetherin, HEK293T cells were seeded in six-well plates and transfected with 4 µg of NL4-3 ΔvpuΔNef IRES eGFP, 1 µg Vpu or Nef expression plasmid and different amounts of tetherin expression vectors (6 well). At two days posttransfection supernatants were harvested and the yield of infectious HIV-1 was determined by a 96-well infection assay on TZM-bl indicator cells.

## Acknowledgments

We thank D. Toso and P. Grob for cryo-EM support, G. Stjepanovic for training and assistance with HDX-MS, and D. Krnavek for technical assistance. EM reconstructions have been deposited in the EMDB and coordinates have been deposited in the RCSB. This research was supported by NIH grants R01 AI 120691 (J. H. H.) and P50 GM082250 (J. H. H.) and an Advanced ERC grant ‘Anti-Virome’ (F. K.) and DFG-funded SFB 1279 and SPP 1923 (F. K.).

## Author Contributions

J.H.H. and X.R. conceived the overall project. K.L.M., C.Z.B., X.R., C.M.S., E.H. performed research. K.L.M. and C.Z.B. performed wild-type and mutant EM analysis respectively, X.R. performed HDX-MS, MS/MS and mutational analysis, C.M.S. and E.H. performed in cell experiments. J.H.H., X.R. and F.K. supervised research. J.H.H., K.L.M., C.Z.B. and X.R. wrote the initial manuscript draft with editing by F.K. All assisted in manuscript preparation.

## Accession numbers

The EM density maps were deposited in the EMDB with accession numbers 7453, 7454 for the dileucine mutant dimer, 7455 for the monomer and 7456, 7457, 7458 for the closed trimer. Particle stacks associated with EMD-7457 and EMD-7458 were respectively deposited to EMPIAR as XXXXX and XXXXX. The whole dataset pre-classification was deposited as EMPIAR-XXXXX. The atomic coordinates for the closed trimer monomeric subunit were deposited with the PDB accession code 6CM9.

## Supplementary Materials

**Table S1.** Plasmids used in this study.

|  |  |
| --- | --- |
| AP-1 core | Hs AP1S3/His-TEVsite-Hs AP1B1 (1-584)/Mm AP1G (1-595)-TEVsite-GST/Mm AP1M1 in pST44 vector |
| AP-1 $\Delta\beta$Arf | AP1B1_I85D/V88D in AP-1 core/pST44 construct |
| AP-1 $\Delta\gamma$Arf | AP1G_L68D/L71E/L102E in AP-1 core/pST44 construct |
| Arf1 | His-TEVsite-Hs Arf1 (17-181)-Q71L in pHis2 vector |
| Tetherin-Nef | His-TEVSite-Hs Tetherin (1-21)- GSDEASEGSG-NL4-3 Nef in pHis2 vector |
| Tetherin-Nef_LL/AA | His-TEVSite-Hs Tetherin (1-21)- GSDEASEGSG-NL4-3 Nef-L164A/L165A in pHis2 vector |
| Tetherin-MRCA Nef | His-TEVSite-Hs Tetherin (1-21)- GSDEASEGSG-O-MRCA Nef in LIC 1B vector |
| MHC-Nef | His-TEVSite-Hs MHC-I (338-365)-NL4-3 Nef in pHis2 vector |
| Nef | His-TEVSite-NL4-3 Nef in pHis2 vector |
| His-MBP-CK1 | His-MBP-TEVsite-Rat CK1δ (1-317) in LIC 2C-T vector |
| HIV-1 ΔVpuΔNef | NL4-3 HIV-1 provirus with deletion of nt3-nt146 in vpu and aa 1,4 and 5 mutated to stop in nef in pBR vector |
| O-MRCA Nef | OMRCA Nef with C-terminal AU1-tag in pCG IRES-eGFP vector |
| NL4-3 Nef | NL4-3 Nef with C-terminal AU1-tag in pCG IRES-eGFP vector |
| O-MRCA Nef C174S | OMRCA Nef C174S with C-terminal AU1-tag in pCG IRES-eGFP vector |
| NL4-3 Nef S169A | NL4-3 Nef S169A with C-terminal AU1-tag in pCG IRES-eGFP vector |
| Vpu | vpu with C-terminal AU1-tag in pCG IRES-eGFP vector |
| Empty vector control | nef with stop codon at position 2,3 and 4 in pCG vector |
| Tetherin | human tetherin in pCG vector |

**Table S2.**
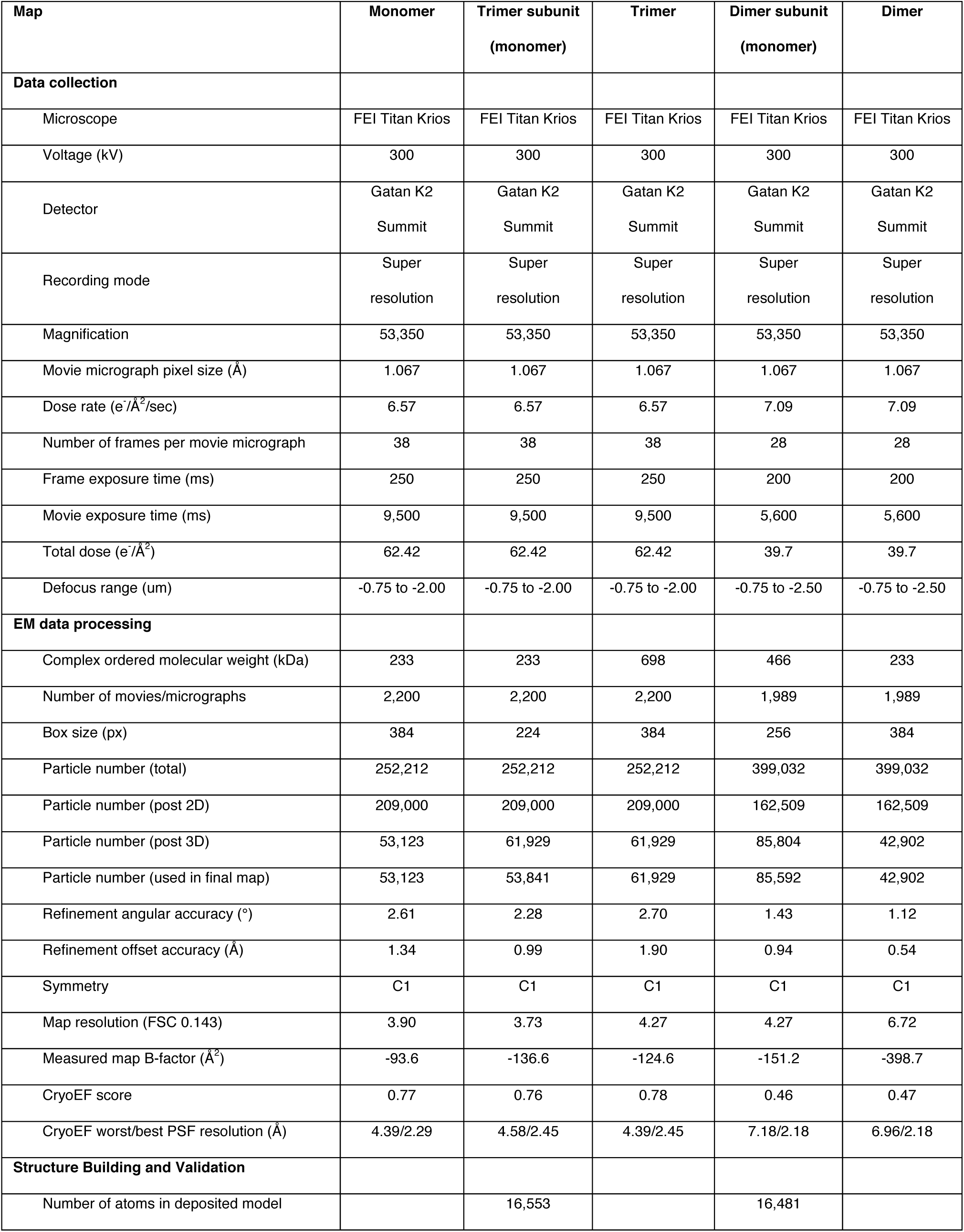
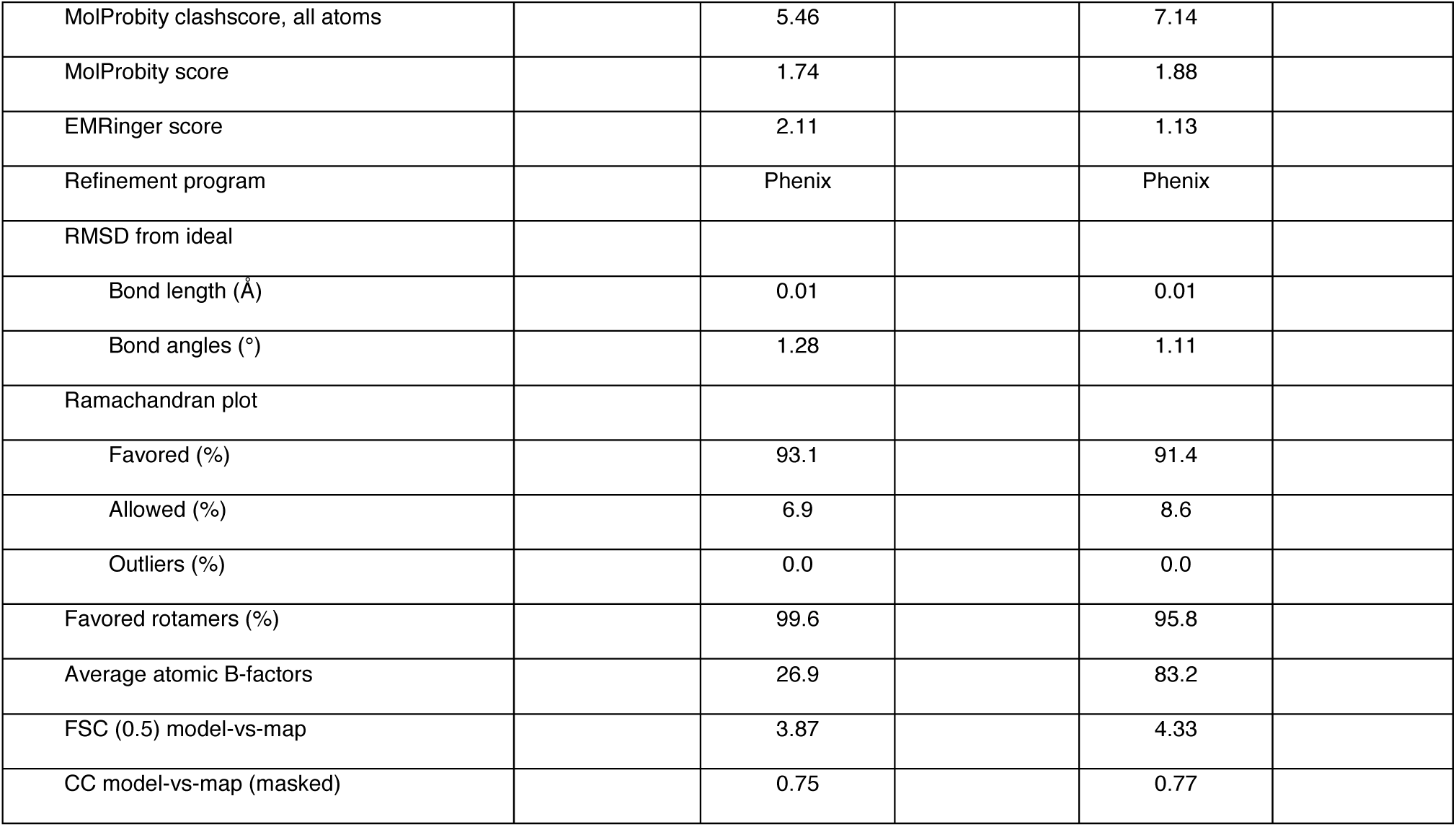
EM and model statistics

**Movie S1.** The closed trimer monomeric subunit reconstruction shown as an assembled AP-1:Arf1:tetherin-Nef complex. The Nef trans AP-1 bridge, the γArf1 trimer, γArf1:AP-1:γ and βArf1:AP-1:β1 interfaces are highlighted. The AP-1 core and γArf1 subunits are sharpened at -50 Å^2^, Nef and βArf1 subunits are unsharpened for clarity.

**Figure S1.**
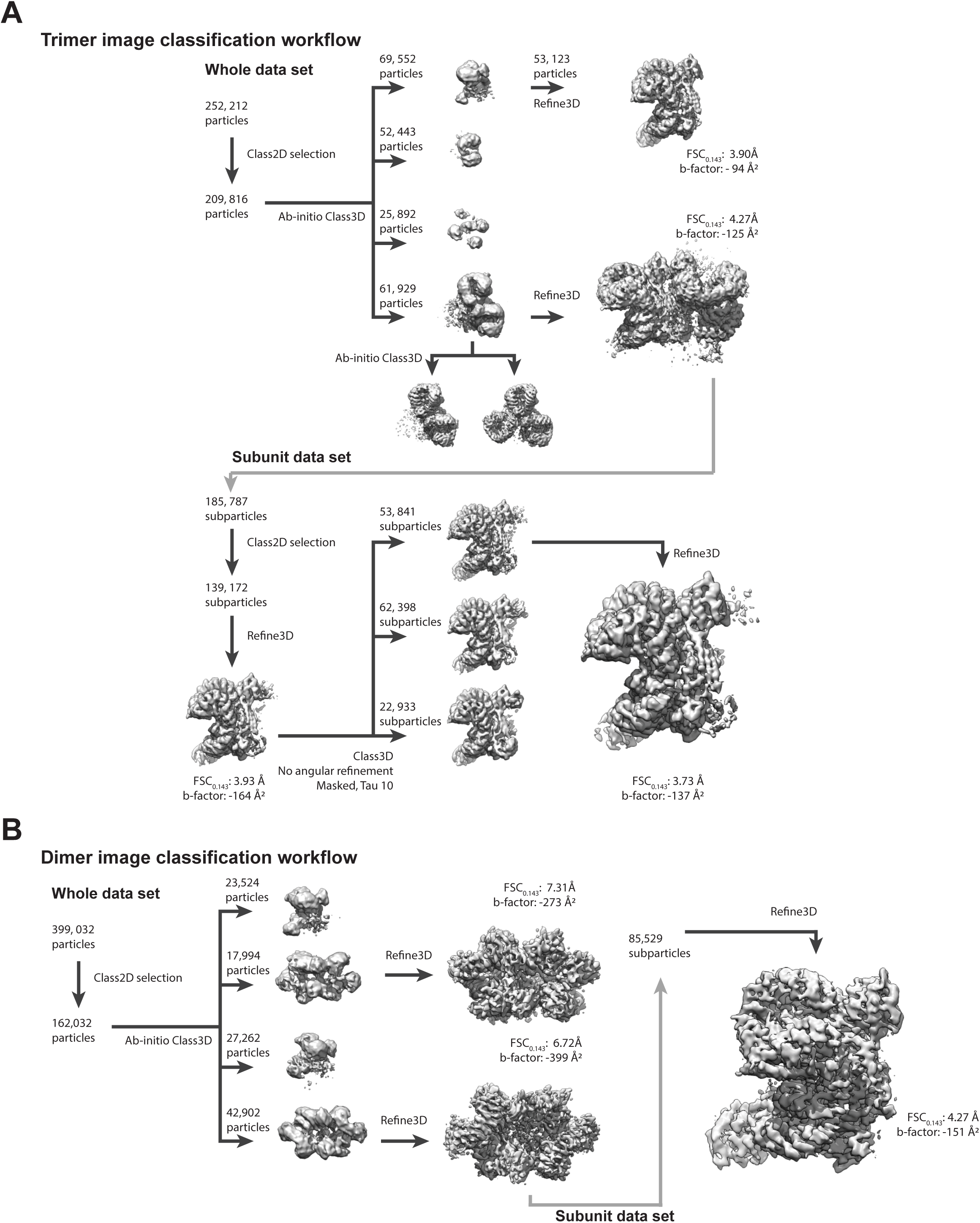
Classification workflow. (A) The wild type closed trimer, monomeric subunit and monomer 3D classification workflow. (B) The Nef dileucine mutant dimer classification workflow. In both cases after cleaning by 2D classification in Relion, initial model generation and 3D classification was performed in cryoSPARC (Punjani et al., 2017). Architecturally homogenous particles had their monomeric subunits extracted (as described in Figure S3) as subparticles for further refinement. Associated with Fig. 1.

**Figure S2.**
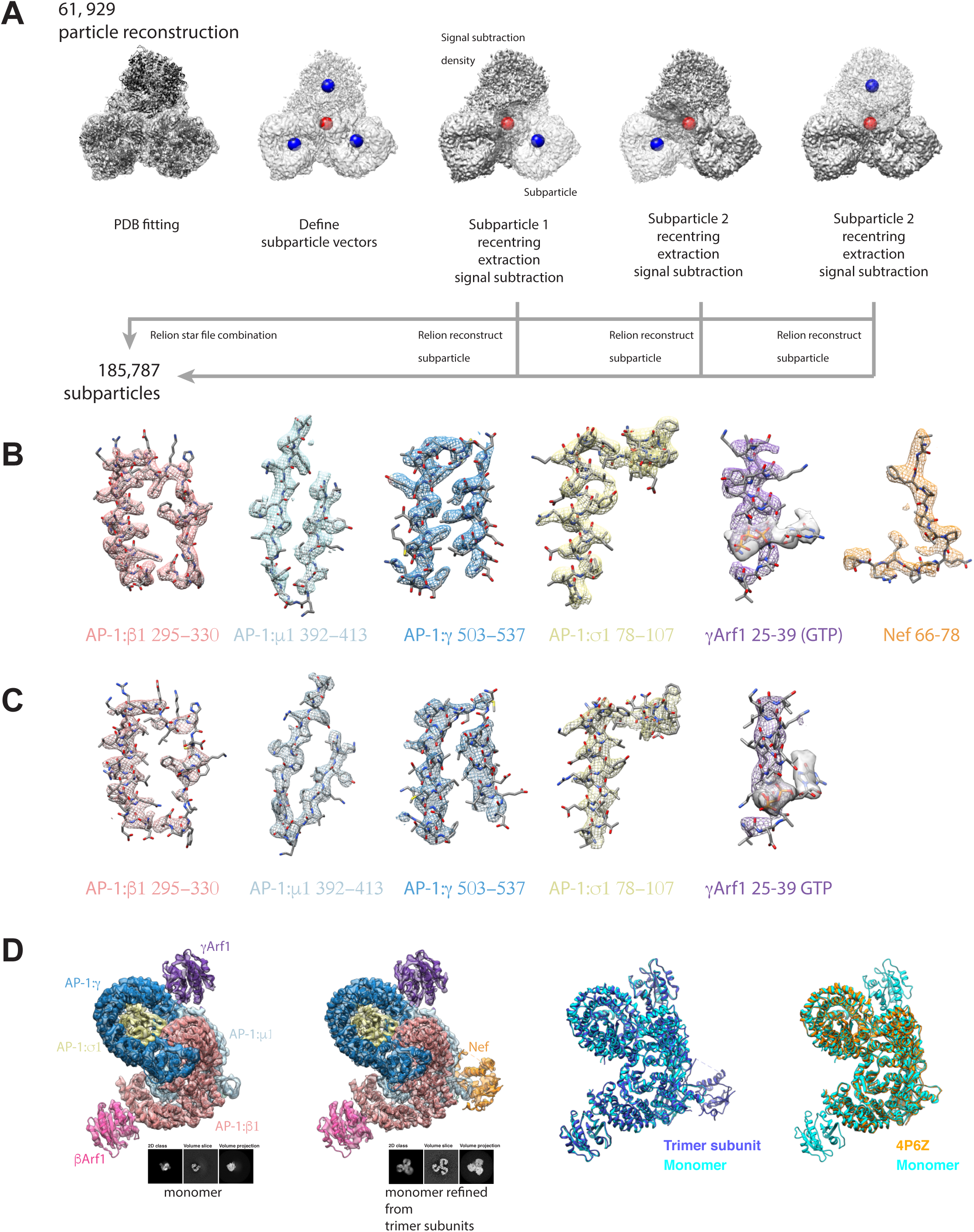
Sub-particle local symmetry expansion and extraction by Relion localized reconstruction. The extraction of monomeric subunits from whole pseudo symmetric complexes (Ilca et al.) as subparticles permits further refinement of a minimal monomeric subunit of the closed trimer. This permits the resolving of high quality structures despite the apparent local variability in trimeric and dimeric oligomeric states. (A) Using a fitted PDB to segment the initial reconstruction masks are created for signal subtraction to leave only the desired subparticle. Vectors describing the sub-particle locations are then used in conjunction with Relion refined parameters to extract and recenter the desired subunit. The subparticle extraction of the closed trimer monomeric subunit increases the particle count from 61,929 to 185,787 particles. (B) The quality of the local subunit density for the closed trimer shown at the measured map B-factor of -137 Å^2^. (C) The quality of the local subunit density for the dileucine mutant dimer shown at the measured map B-factor of -151 Å^2^. (D) Monomers were identified in the AP-1:Arf1:Tetherin-Nef sample having both γArf1 and βArf1 bound but no resolvable Nef core. The monomeric subunit of the trimer was determined and has a resolvable Nef core. The AP-1 core and γArf1 subunits are sharpened at -50 Å^2^, Nef and βArf1 subunits are unsharpened for clarity. Both structures have the same AP-1 heterotetrameric core conformation. Thus both the AP-1 core of the monomer and monomeric subunit of the closed trimer are found to be hyper-unlocked when compared to the hyper-unlocked structure of a tetherin-Vpu fusion AP-1 structure (4P6Z) (Jia et al., 2014). Associated with Fig. 1.

**Figure S3.**
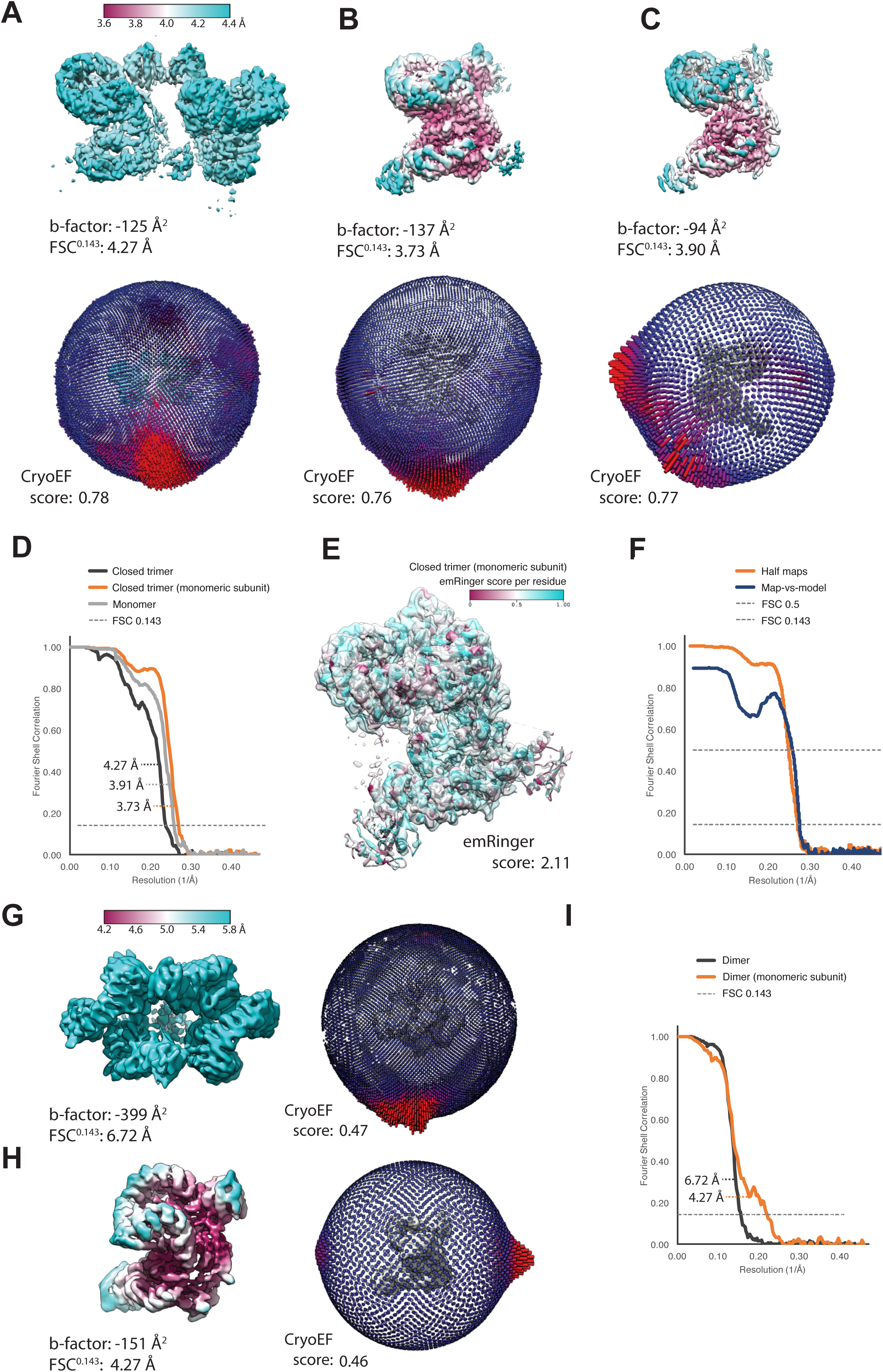
Map and model validation. (A) Local resolution estimation of the whole trimer complex, (B) the closed trimer monomeric subunit refinement and (C) the monomer refinement with angular distributions of the final refinements and cryoEF scores. (D) Whole reconstruction gold-standard FSC resolution estimation of the closed trimer compared to the trimer monomeric subunit and true monomer. (E) emRinger and (F) map-vs-model validation of the closed trimeri monomeric subunit structure. (G) Local resolution and angular distribution for the dileucine mutant dimer and (H) dimer subunit. (I) Whole reconstruction gold-standard FSC of the dileucine dimer. Associated with Fig. 1.

**Figure S4.**
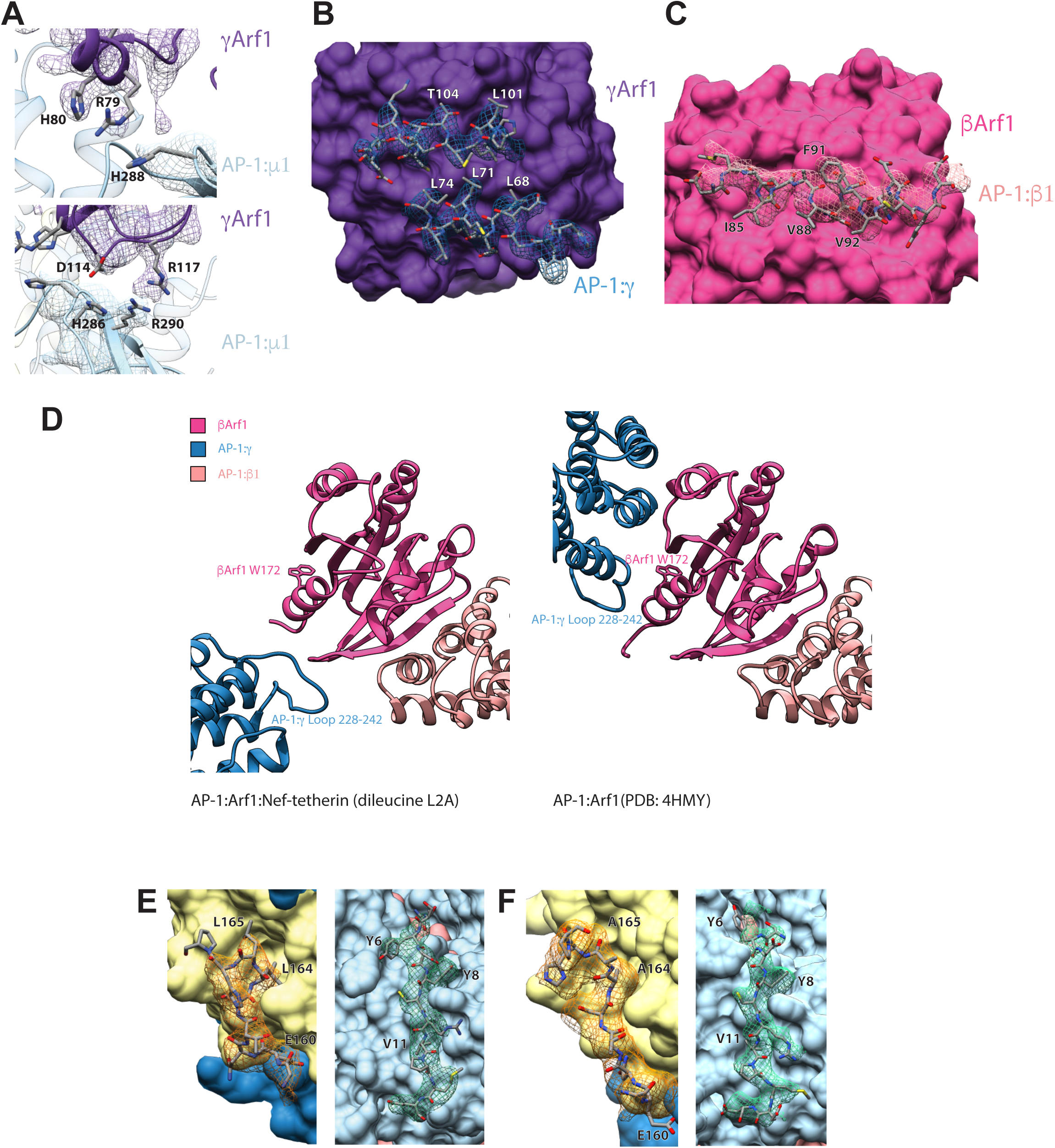
Arf1 architecture and binding interfaces. (A) γArf1 contacts to AP-1:μ1 as well as (B) AP-1:γ in the closed trimer. (C) βArf1 contacts to AP-1:β1 in the closed trimer. (D) In the dileucine mutant dimer a novel βArf1:AP-1:γ interface is present (left) when compared to the crystal structure βArf1:AP-1:γ dimer contact in 4HMY (right). Both AP-1:σ1 and μ1 are cargo binding sites are occupied by the LL motif of Nef and YxxΦ of tetherin respectively in (E) the wild-type monomer and (F) dileucine mutant dimer. Associated with Fig. 1.

**Figure S5.**
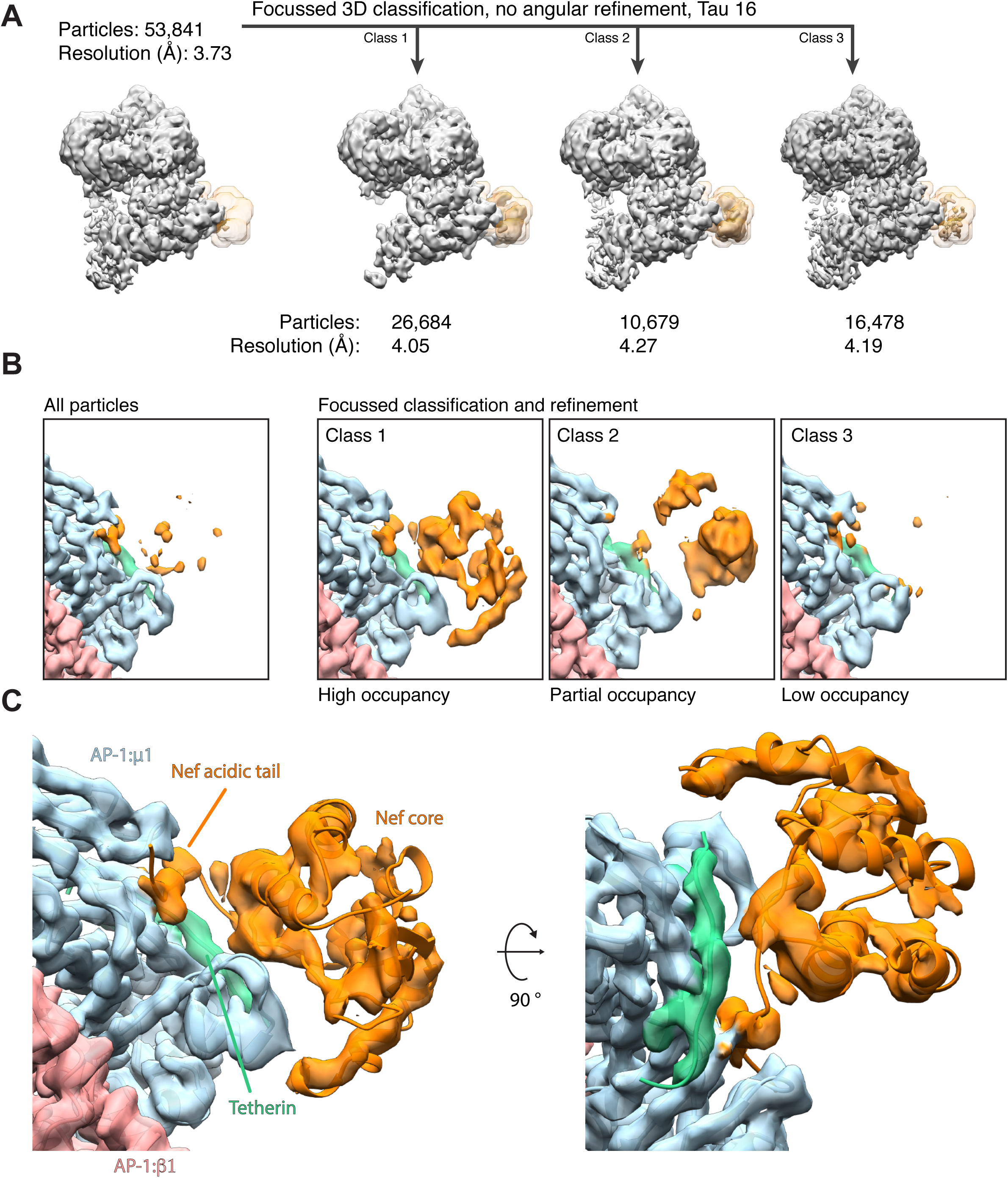
Resolving Nef in the closed trimer by focused classification. (A) A mask (transparent orange) was used to enclose the segmented density of Nef and focused 3D classification without angular sampling determined 3 classes that could be refined to between 4.06 – 4.19 Å. (B) The three structures represent reveal a high, partial and low occupancy state for Nef in the closed trimer monomeric structure. (C) The high occupancy state shows a very good agreement between the Nef model and the refined density after focused classification. All volumes are B-factor sharpened by -50 Å^2^. Associated with Fig. 1.

**Figure S6.**
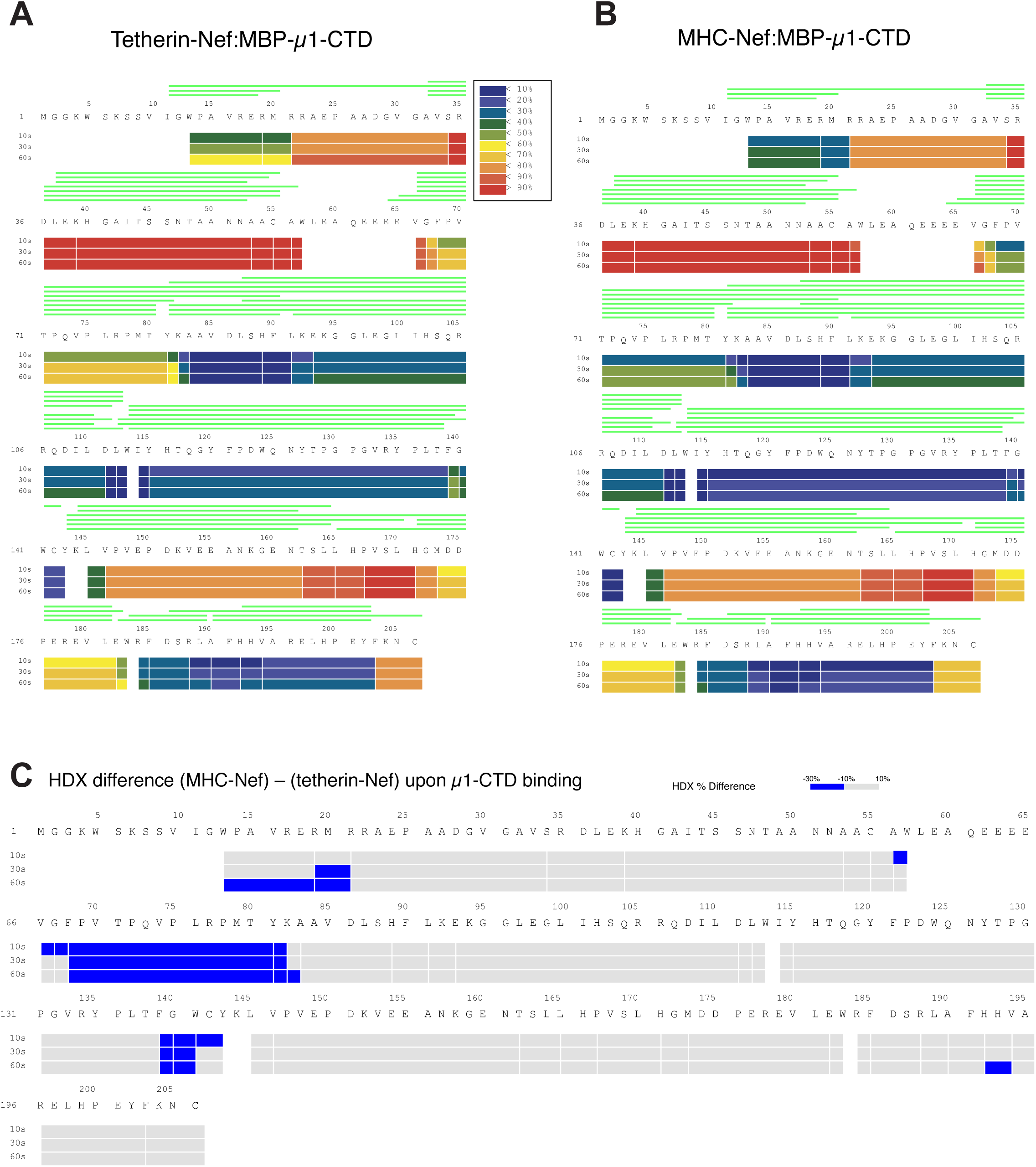
Supporting data for HDX data. Deuterium exchange data for Nef when bound to MBP-tagged AP-1: μ1-CTD in the case of (A) tetherin cargo or (B) MHC-I cargo fusion. HDX-MS data are shown in heat map format where peptide coverage is represented as rectangular strips above the protein sequence. (C) The difference map between bound MHC-I-Nef and tetherin-Nef reveals greater areas of protection on Nef when bound with MHC-I cargo. Associated with Fig. 4.

